# Mechanisms of microtubule dynamics from single-molecule measurements

**DOI:** 10.1101/2025.06.25.661545

**Authors:** Saradamoni Mondal, Eric Bonventre, William O. Hancock, Luke M. Rice

**Affiliations:** Department of Biomedical Engineering, The Pennsylvania State University, University Park, PA 16802; Department of Biophysics, University of Texas Southwestern Medical Center, Dallas, TX 75390; Department of Chemistry, The Pennsylvania State University, University Park, PA 16802; Department of Biochemistry, University of Texas Southwestern Medical Center, Dallas, TX 75390

## Abstract

Microtubules are dynamic polymers of αβ-tubulin heterodimers that organize the intracellular space and mediate faithful chromosome segregation. Microtubule function depends on dynamic instability, the apparently random GTPase-dependent switching between growing and shrinking. Microtubule dynamics derive from biochemical properties of individual tubulin subunits and how they interact with the polymer end, a complex environment where individual tubulins can have different numbers of neighbor contacts. A fundamental understanding of microtubule dynamics has been difficult to establish because of challenges measuring the number, strength, and nucleotide-dependence of tubulin binding sites on the microtubule end. We used an improved single-molecule assay to measure tubulin:microtubule interactions. In addition to the two expected classes of binding site (longitudinal and corner), we identified previously unrecognized third binding interaction. We further show that nucleotide state strongly influences the strength of inter-protofilament contacts, with little effect on intra-protofilament contacts, and that a mutation can modulate this nucleotide effect. By uncovering a new tubulin binding state on the microtubule end, clarifying how GDP influences microtubule stability, and demonstrating that the nucleotide effects are allosteric and tunable, these single-molecule measurements and accompanying computational simulations provide rich new biochemical insight into the fundamental mechanisms of microtubule dynamics.

## INTRODUCTION

Microtubules are dynamic cytoskeletal polymers of αβ-tubulin that orchestrate chromosome segregation, provide tracks for motor-based trafficking, and underlie cellular mechanics. Microtubule function depends on dynamic instability (Mitchison & Kirschner, 1984), the GTPase-dependent switching of individual polymers between phases of growing and shrinking. The rates of microtubule growing or shrinking reflect the net association or dissociation of αβ-tubulins, which are dictated by the biochemistry of αβ-tubulin:microtubule interactions (Brouhard & Rice, 2018; Cleary & Hancock, 2021; Gudimchuk & McIntosh, 2021; Howard & Hyman, 2003). The understanding of microtubule growing, shrinking, and switching has improved remarkably in recent years, but there are still fundamental unresolved questions about the molecular mechanisms underlying dynamic instability.

Two main obstacles explain why it has been difficult to define biochemical mechanisms of microtubule dynamics. First, microtubule ends are biochemically and configurationally heterogeneous environments presenting a poorly defined mix of binding sites that differ in strength owing to different numbers of longitudinal and lateral interactions (Atherton et al., 2017; Chretien et al., 1995; McIntosh et al., 2018; VanBuren et al., 2002). Second, there is an inference problem: it has generally not been possible to observe microtubule growing and shrinking at the level of individual αβ-tubulins, so estimates of the biochemical rate constants for tubulin:microtubule interactions have instead been obtained indirectly by fitting models to observed microtubule growing and shrinking rates. However, this inference process can produce degenerate solutions which can lead to conflicting predictions (Cleary et al., 2022; Coombes et al., 2013; Gardner et al., 2011; Gudimchuk et al., 2020; Margolin et al., 2012; VanBuren et al., 2002; Zakharov et al., 2015).

Recently introduced measurements and analyses have shown promise to enable a deeper and more comprehensive understanding of the mechanisms of microtubule dynamics. We showed that simultaneous analysis of microtubule growth rates and fluctuations narrowed the degeneracy of possible solutions, revealing that αβ-tubulins associate more slowly with the microtubule end than had previously been thought (Cleary et al., 2022). We also used interferometric scattering (iSCAT) microscopy combined with nanogold labeled αβ-tubulin to measure individual tubulin associations with and dissociations from growing microtubule ends (Mickolajczyk et al., 2019). This more direct binding study overcame disadvantages associated with inference, but it had its own limitations: the focus on growing microtubules required unlabeled tubulin, and the influence of this ‘invisible’ tubulin made it hard to directly interpret single-molecule observations in terms of biochemical off-rates.

In the present study, we used an improved single-molecule assay that eliminates unlabeled tubulin, obtaining new data that more faithfully report on the biochemistry of tubulin:microtubule interactions. Our measurements revealed three classes of tubulin binding site on the microtubule end that we ascribe to a weak single longitudinal contact, a strong longitudinal + lateral (corner) contact, and an unexpected interaction of intermediate affinity. The measured residence times for tubulin:microtubule binding were readily interpretable in terms of dissociation rates and recapitulated measured microtubule growth rates when used in a computational model for microtubule dynamics. Measurements in the presence of GDP revealed that nucleotide state predominantly affects the strength of lateral tubulin:tubulin contacts, with only modest effects on longitudinal interactions. Finally, we show that a tubulin mutation that hyperstabilizes microtubules does so by attenuating the allosteric response to GDP. By revealing a new biochemical state in microtubule elongation, defining how GDP affects tubulin:microtubule interactions, and showing that the response to GDP is ‘tunable’, our findings provide a rich set of new insights into the biochemical mechanisms of microtubule dynamics.

## RESULTS

### An improved single-molecule assay more reliably measures tubulin:microtubule interactions

We previously used single-molecule measurements of nanogold-labeled yeast tubulin interacting with a growing microtubule end to detect and quantify two classes of binding sites that we ascribed to longitudinal and ‘corner’ (longitudinal+lateral) interactions (Mickolajczyk et al., 2019). In principle, characteristic dwell times from such an experiment should be concentration-independent because they should be inversely related to the off-rate constants for the available binding sites. However, the measured characteristic dwell times decreased with increasing concentrations of unlabeled tubulin, which indicated that the unlabeled tubulin was affecting the behavior of the labeled tubulin. Consequently, the dwell time distributions could not be unambiguously interpreted in terms of binding affinity or off-rate.

To improve the biochemical fidelity of our single-molecule assay for tubulin:microtubule interactions, we eliminated the unlabeled tubulin that previously complicated interpretation of the measured dwell times. We also switched from recombinant yeast tubulin to recombinant human tubulin. In this refined assay (Fig. 1A), we use iSCAT (interferometric scattering) microscopy (see Methods) to detect individual nanogold-labeled tubulins reversibly binding the plus- and minus-ends of static GMPCPP-stabilized microtubules, with high contrast and at 125 frames per second. Using this approach, we observed dwell times spanning 5 orders of magnitude (milliseconds to 100s of seconds), with notably longer dwell times than in our prior work (Mickolajczyk et al., 2019).

**Fig. 1.**
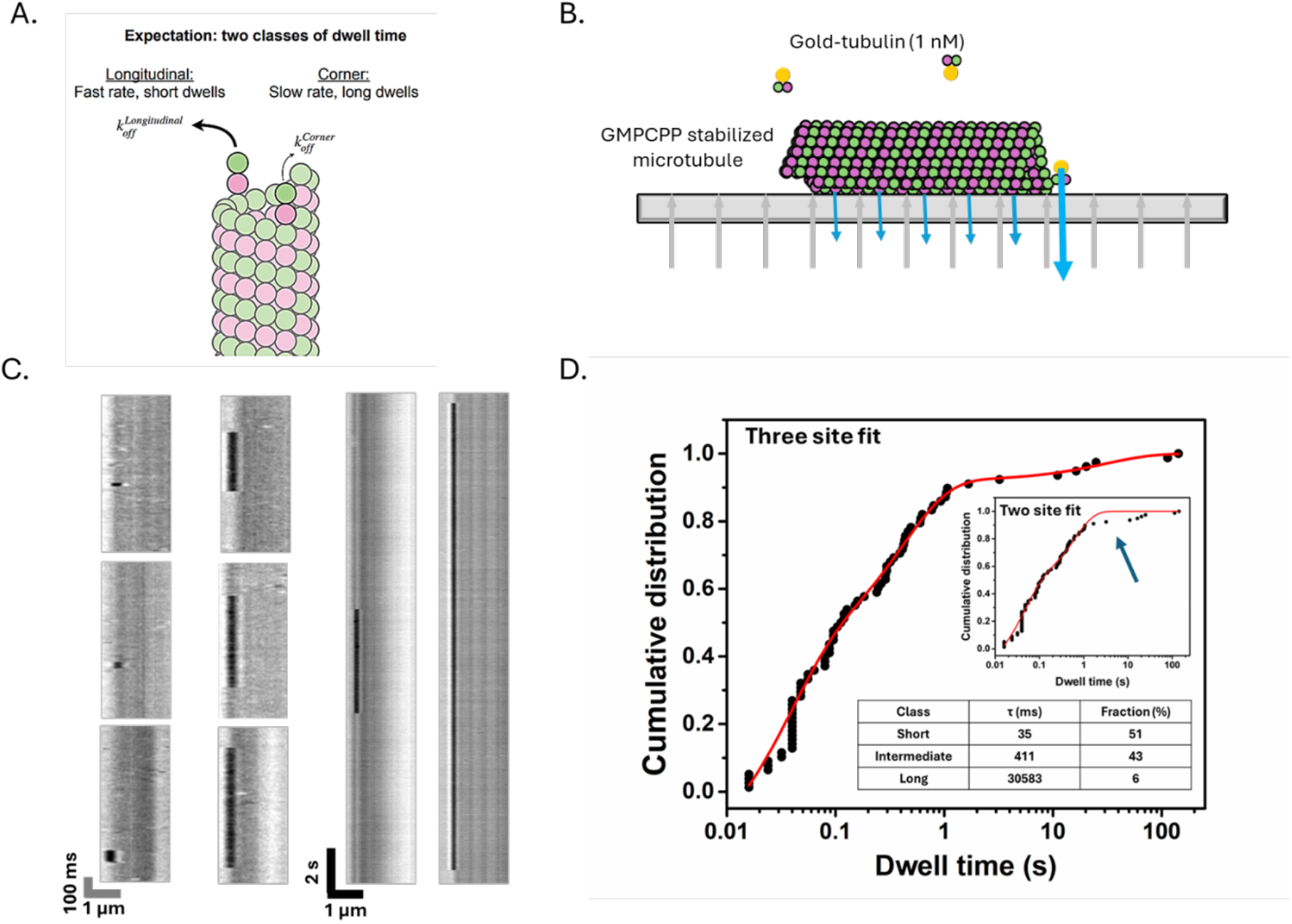
Single-molecule measurements of gold-labeled recombinant human tubulin (gold-tubulin) reversibly interacting with microtubule ends. (A) Diagram of microtubule plus-end showing longitudinal and corner interactions (where k_off_ is inverse of dwell time). (B) Schematic diagram of the single-molecule assay. Stable microtubule seeds were attached to coverslips using an anti-DNP antibody, incubated with 1 nM gold-tubulin in 1 mM GMPCPP, and imaged using iSCAT microscopy. (C) Representative kymographs showing reversible binding events with different residence times. (D) Cumulative distribution function (CDF) for the measured dwell times (circles) with the tri-exponential fit (red line); n =78 dwell events. The best fit characteristic times are 35 [26-43] ms representing 52 % of all dwells, 411 [334-488] ms representing 42 % of all dwells, and 31 [0-62] s representing 6 % of all dwells; brackets give 95% confidence intervals. Inset: CDF of measured dwell time distribution fit with a bi-exponential function that fails to recapitulate the long dwells (arrow).

### Dwell time distributions reveal three classes of binding site on stabilized microtubule ends

Based on the expected ratio of longitudinal to corner sites on the end of a stabilized 13 protofilament microtubule (Chaaban et al., 2018; Tilney et al., 1973), we expected to observe two classes of dwell times: a short class reflecting a fast off-rate for lower affinity longitudinal interactions and a long class reflecting a slow off-rate for higher affinity corner interactions, in roughly a 12:1 ratio (Fig. 1A). However, when we attempted to fit the measured dwell times with a bi-exponential function representing two classes of binding site (dwell time), the longest dwells were systematically unaccounted for at both ends of the microtubule (Fig. 1D, inset; Fig. S1; see Methods). Thus, the measurements must report on more than two classes of tubulin binding site. A tri-exponential function (see Methods) captured the full range of measured dwells, yielding characteristic plus-end dwell times of 34.7 ms (“fast” events, 52% of total dwells), 411 ms (“intermediate” events, 42% of total dwells), and 30.5 s (“slow” events, 6% of total dwells) (Fig. 1D; see Fig. S1 for minus-end results). Thus, our improved assay unexpectedly revealed three distinct classes of binding site on the microtubule end.

### Biochemical interpretation of the three binding sites: one corner and two distinct longitudinal interactions

We assign the short (35 ms) class of dwells to pure longitudinal interactions for three reasons: their timescale is comparable to the 50 ms dwells previously ascribed to longitudinal interactions (Mickolajczyk et al., 2019), the timescale is consistent with a relatively weak interaction, and their proportion (∼50% of all events) comports with the larger number of longitudinal vs corner sites expected on the stabilized microtubule end. Likewise, we assign the long class of dwells to corner interactions: the 30 s characteristic dwell time is consistent with a high affinity interaction (the derived off-rate of 0.03 s^-1^ converts to 30 nM affinity assuming a typical on-rate constant of 1 μM^-1^s^-1^ (Northrup & Erickson, 1992); see next section), and their proportion (∼6 % of all events) is close to the expected fraction of corner binding sites (1/13) on the stabilized microtubule end. The long dwells we report here are ∼15-fold longer than those we measured previously (Mickolajczyk et al., 2019), validating the decision to eliminate unlabeled tubulin from the assay.

What kind of interaction do the intermediate dwells reflect? In principle they might be an artifact of some gold particles containing more than one tubulin and consequently able to make additional microtubule contacts, but we discard this possibility because three-phase binding persisted at both microtubule ends when we used a 10-fold lower gold:tubulin ratio (Fig. S2). The intermediate dwells also seem too numerous (∼40% of dwells on top of the 6% already assigned as bona fide corners) and too close in timescale (within 10-fold) to the short dwells to represent an alternative corner interaction. We argue instead that the simplest model to explain the intermediate dwells is that they represent a second, previously undetected class of higher-affinity longitudinal interaction at the microtubule end. We imagine that upon binding the end of a protofilament, tubulin initially forms a weak longitudinal contact that subsequently ‘matures’ through some kind of isomerization (conformational change) to make a ∼10-fold stronger interaction.

### Using biochemical off-rates derived from measured dwell times, a minimal kinetic model that includes the isomerization reaction can recapitulate microtubule growth rates

To test if a model including this new binding site can recapitulate microtubule growth rates, we incorporated into a kinetic model for microtubule polymerization (Cleary et al., 2022; Kim & Rice, 2019; McCormick et al., 2024; Mickolajczyk et al., 2019; Piedra et al., 2016) an isomerization reaction that results in a higher affinity longitudinal interaction. For simplicity, we consider the isomerization to be irreversible. The model simulates microtubule dynamics one biochemical event (association, dissociation, and now isomerization) at a time, effectively generating a molecular movie of how microtubules grow. A tubulin that binds longitudinally to the end of a protofilament will do one of three things: dissociate rapidly according to the weaker longitudinal affinity; isomerize to the stronger, more slowly dissociating longitudinal interaction; or be ‘promoted’ to a corner interaction by association of another tubulin. Thus, the parameters defining the reaction kinetics in the model (Fig. 2B) are: an association rate constant k_on_ that determines how fast tubulins associate with protofilament ends, three off-rate constants that specify how quickly tubulins dissociate from different kinds of binding site (weak longitudinal, stronger longitudinal, and corner), and an isomerization rate constant that sets the rate of a tubulin ‘maturing’ from weak to stronger longitudinal states.

**Fig. 2.**
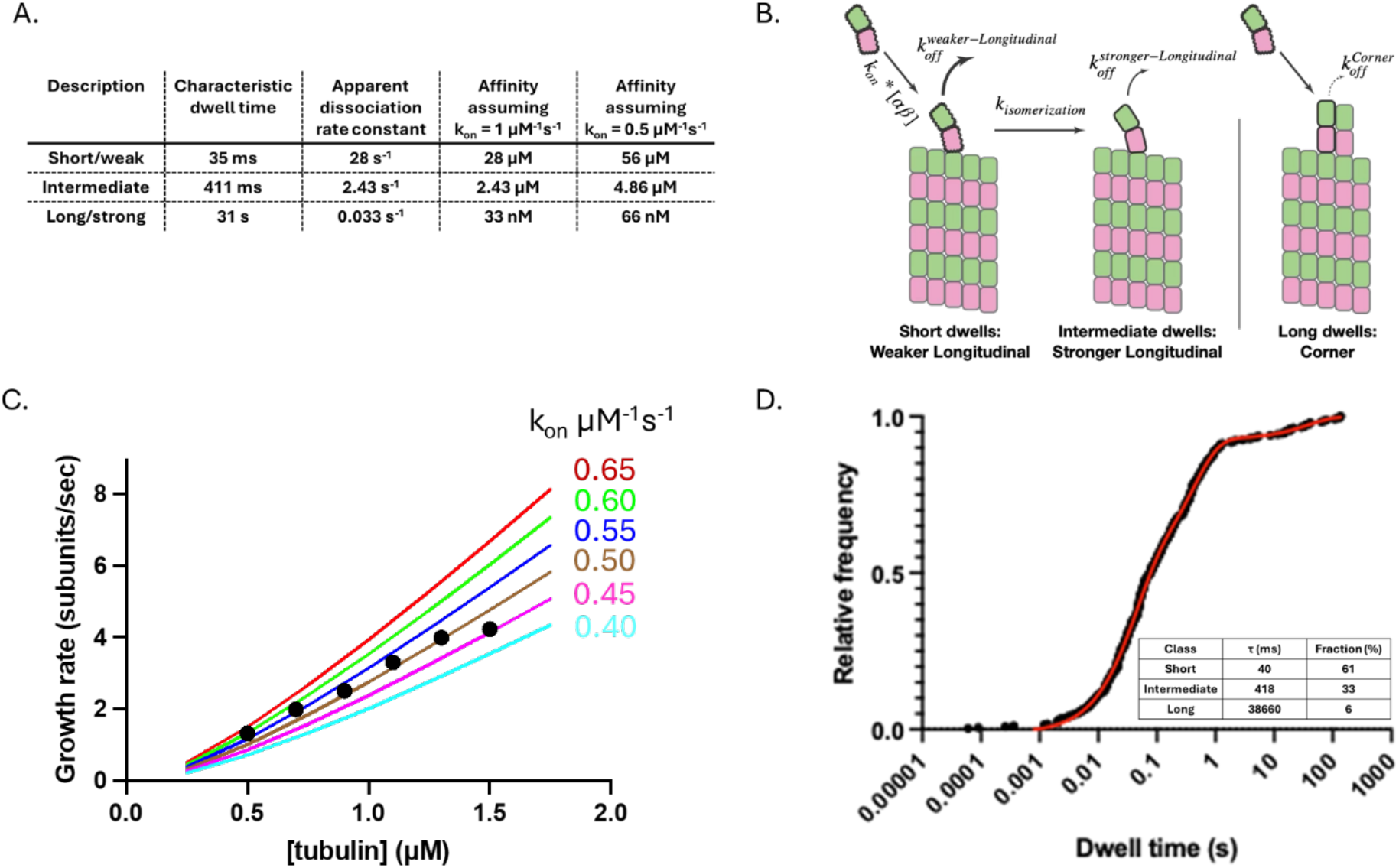
Kinetic modeling of microtubule polymerization incorporating isomerization to a ‘matured’ longitudinal interaction. (A) Summary of measured characteristic dwell times, the calculated dissociation rate constants, and the corresponding affinities under different assumptions about k_on_. (B) Schematic representation of the proposed model where a tubulin dimer initially binds with weak affinity and can undergo a conformational change to a higher-affinity “locked-in” state. (C) Schematic diagram illustrates the kinetic model, including association, dissociation, and isomerization steps. (D) Simulated microtubule growth rates at various k_on_ values compared to experimental data for GMPCPP-tubulin polymerization. (E) CDF of simulated tubulin dwell times, which closely resembles the experimentally observed dwell times.

In principle, all model parameters except k_on_ should be specified by the three-phase fit to the dwell time distribution (see Methods): the characteristic dwell times specify the off-rate constants for the corner and two types of longitudinal interactions, and the isomerization rate can be derived from the proportions of fast and intermediate dwells. Indeed, the relative fraction of fast and intermediate binding events is related to the relative magnitudes of the off-rate from the weak longitudinal state and the isomerization rate (see Methods). Thus, and taking as a priori information the dissociation and isomerization rates determined from the analysis of dwell times, we sought to test if there was a reasonable value for k_on_ that could recapitulate measured microtubule growth rates. This parsimonious approach is highly constrained and, if successful, should yield a model that simultaneously matches microtubule growth rates and the measured dwell times.

We simulated microtubule growth rates for a range of assumed association rate constant k_on_ and compared the resulting predictions to concentration-dependent microtubule growth rates measured previously (Cleary et al., 2022) (Fig. 2C). Simulated growth rates (lines in Fig. 2C) increase as the assumed value of k_on_ increases, and the best match to measured growth rates occurred for k_on_ = 0.5 μM^-1^s^-1^, a value close to the one we estimated previously from joint analysis of microtubule growth rates and fluctuations (Cleary et al., 2022). As expected, dwell time distributions extracted from the simulations showed three phases, and the amplitudes and characteristic dwell times of these phases closely matched the experimental data that were used as constraints (Fig. 2D).

### Three classes of binding site can also be observed on growing microtubule ends

Growing microtubules have different end configurations and may present different conformations of tubulin compared to stabilized microtubule ends (Atherton et al., 2017; Chaaban et al., 2018; Chretien et al., 1995; McIntosh et al., 2018; van den Berg et al., 2023). To test whether the three classes of interaction we observed on stabilized microtubule ends are also present on growing microtubule ends, we measured single-molecule dwell times using growing microtubules as the substrate. At a low concentration of unlabeled tubulin (0.25 µM), the measured dwell time distribution clearly showed three-phase behavior (Fig. 3) with characteristic dwell times of 74 ms (fast events, 62%), 573 ms (intermediate events, 33%), and 3.5 s (slow events, 5%). Fast and intermediate dwells times on the growing microtubule end differed only slightly from those measured on the stabilized microtubule end, whereas the slow dwell time on the growing microtubule end was ∼10-fold shorter than what we measured in the absence of unlabeled tubulin. This selective reduction of the slow dwell times can be explained by preferential trapping of longer-dwelling gold-tubulins by unlabeled tubulin and is consistent with the observation that 11% of the binding events were irreversible (Fig. 3C). At a higher concentration of unlabeled tubulin (1 µM), three phase behavior was no longer evident: two characteristic times of 88 ms (fast events, 58%) and 2 s (42%) were sufficient to recapitulate the dwell time distribution. This slower dwell time likely reflects a mix of intermediate dwells and the shortest remaining reversible corner events, with the 22% of irreversible binding events representing trapping of longer binding events into the growing microtubule lattice. In summary, the ‘matured’ longitudinal interaction can also be detected on a growing microtubule but is obscured if the concentration of unlabeled tubulin is too high.

**Fig. 3.**
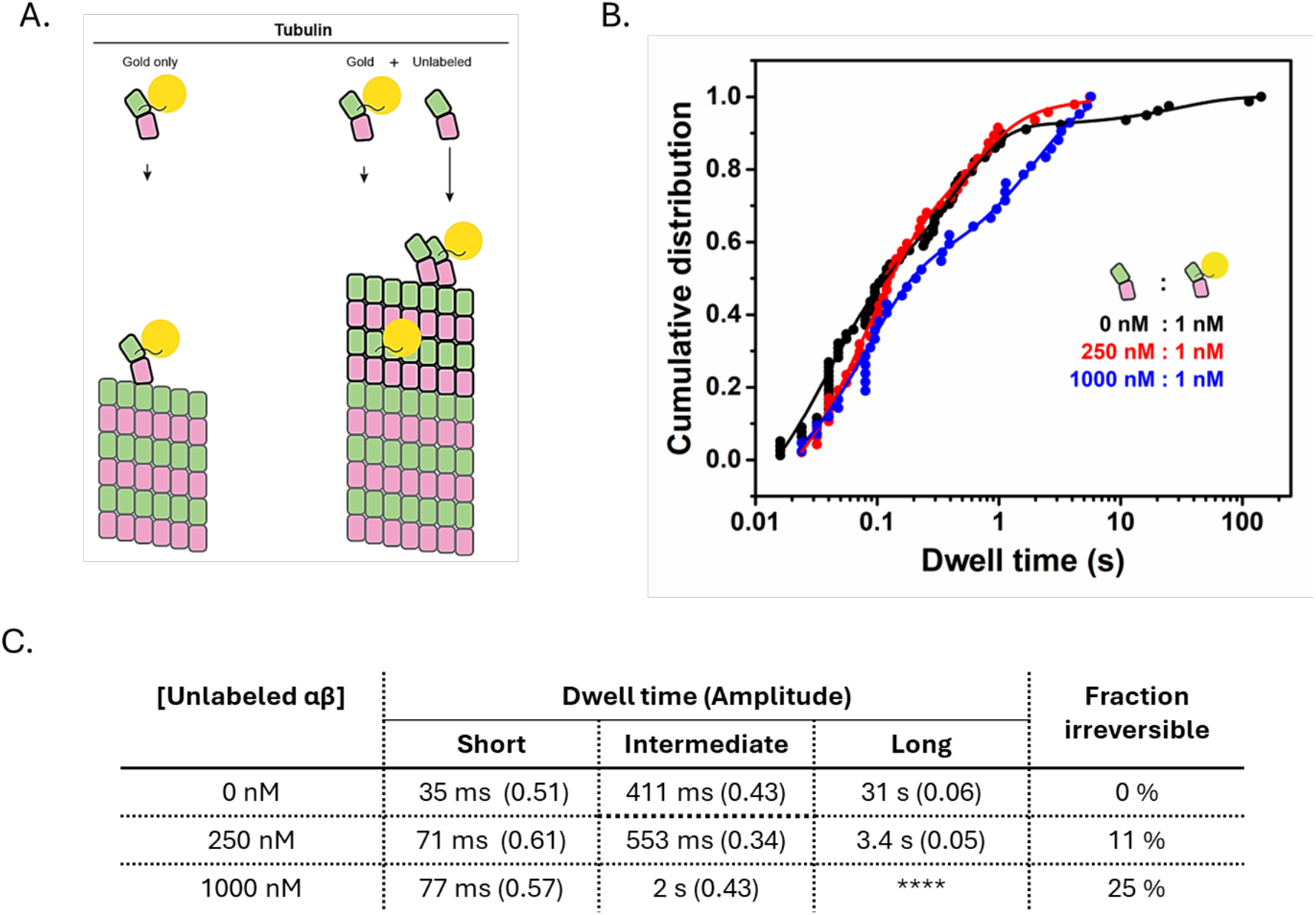
The matured longitudinal interaction is also detectable on a growing microtubule end. (A) Cartoon of tubulin binding and incorporation into the microtubule lattice at increasing concentrations of unlabeled tubulin. Left: without unlabeled tubulin, trapping of gold tubulin does not occur (all associations are reversible). Right: trapping of gold-tubulin becomes more likely as the concentration of unlabeled tubulin increases. (B) Cumulative distribution of tubulin dwell times without unlabeled tubulin (black; data reproduced from Fig. 1), and with 0.25 µM (blue, n=45), and 1 µM (red, n=55) unlabeled tubulin. For 0.25 µM unlabeled tubulin, the best fit characteristic dwell times are 71 [55-86] ms, 553 [170-940] ms and 3.4 [0-14] s. For 1.0 µM unlabeled tubulin, the best-fit characteristic times are 77 [66-89] ms and 2 [1.8-2.2] s. (C) Table summarizing the characteristic dwell times and fraction of irreversible events.

### GDP significantly weakens lateral tubulin:tubulin contacts, with little effect on longitudinal contacts

GDP destabilizes tubulin:tubulin interactions compared to GTP, but the specific mechanism remains unresolved (Alushin et al., 2014; Fedorov et al., 2019; Hemmat & Odde, 2021; Igaev & Grubmuller, 2020, 2022; Manka & Moores, 2018; McCormick et al., 2024; Piedra et al., 2016; Tong & Voth, 2020; Wang & Nogales, 2005; Zhou et al., 2023). To directly assess how nucleotide state impacts the affinity of tubulin:microtubule interactions, we measured dwell times in the presence of GDP (see Methods). Contrary to the experiments in GMPCPP, and despite a similar event frequency (Fig. 4A), we did not observe long dwells in GDP: no dwells exceeded 3.5 s (Fig. 4A). A biexponential function was sufficient to fit the distribution of dwells at the plus end, yielding characteristic times of 28 ms (fast events, 70%) and 232 ms (intermediate events, 30%). Both characteristic times are approximately twofold shorter than corresponding times we measured in GMPCPP. Based on the similar proportions, the fast and intermediate events observed in GDP likely correspond to the weak and stronger forms of longitudinal interaction. The minus end yielded similar results: no long dwells, and only two characteristic dwell times of 45 ms (fast, 78%) and 153 ms (intermediate, 22%) were needed to fit the distribution (Fig. S3). Thus, GDP modestly weakens longitudinal tubulin:microtubule interactions (∼twofold lower affinity), but it attenuates lateral interactions by at least 100-fold (estimated based on the reduction required to make a 30.6 s dwell time in GMPCPP indistinguishable from the 153 ms intermediate dwell time in GDP). In principle, the reduced corner affinity in GDP could result from either a weakening of the lateral tubulin:tubulin interface or a stiffening of the incoming tubulin dimer that increases the ‘straightening penalty’ inherent in lattice incorporation (Brouhard & Rice, 2018).

**Fig. 4.**
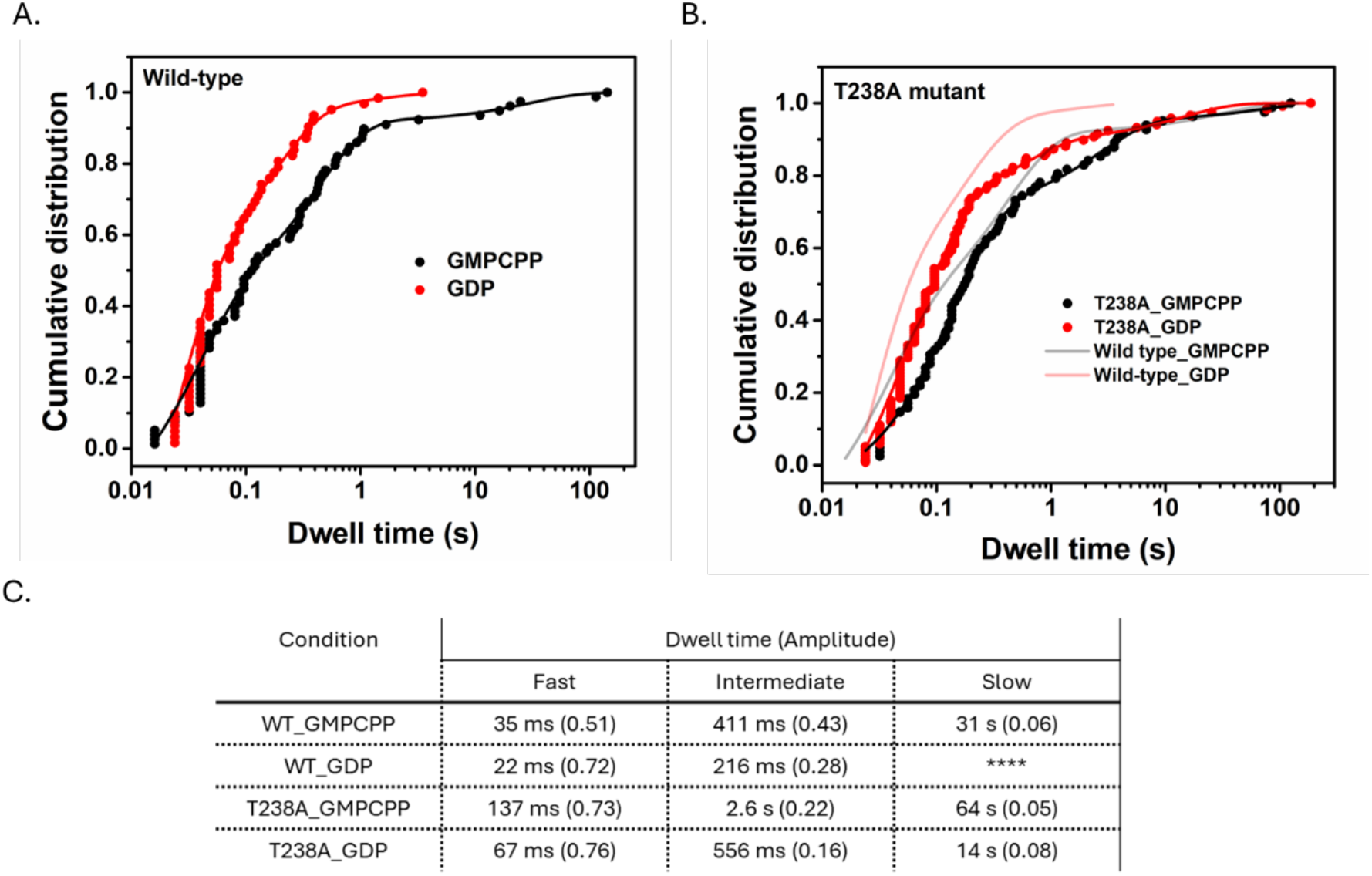
Dwell time distributions for wild-type and mutant tubulin in GMPCPP and GDP conditions. Cumulative distribution of dwell times for wild-type (A) and mutant tubulin (B) fitted with multi-exponential decay functions. In both panels, the black data points and the fitted curves represent GMPCPP, while the red data points and curves correspond to GDP. For GDP-bound wild-type tubulin, the data were best fitted with a bi-exponential function (n =62 dwell events), yielding characteristic times: 22 [19-27] ms and 217 [190-240] ms; whereas the mutant tubulin required a tri-exponential fit under both GDP and GMPCPP conditions. (n =118 and 85 dwell events). For GMPCPP-bound mutant tubulin, the best-fit characteristic times are 137 [130-145] ms, 2.6 [2.0-3.4] s and 67 [0-140] s; in GDP, the best fit characteristic times are 67 [58-71] ms, 556 [310-800] ms, and 14 [7.2-21] s. Table below each graph summarize the binding events, categorized as fast, intermediate, and slow phases, with corresponding dwell time and fraction of contributions (in parentheses).

### A mutation that hyperstabilizes microtubules counteracts the weakening effects of GDP

An increasing number of mutations are known to alter microtubule dynamics (Geyer et al., 2015; Gupta et al., 2002; Hoff et al., 2022; Li et al., 2020; Macaluso et al., 2025; Machin et al., 1995; Park et al., 2021; Roostalu et al., 2020; Sage et al., 1995; Ti et al., 2016), but as with nucleotide state, the underlying biochemical mechanism is often difficult to decipher. We used our single-molecule assay to determine how β:T238A, a well-studied buried mutation that can reduce microtubule shrinking rates by up to 100-fold (Geyer et al., 2015; Ye et al., 2020), increases microtubule stability.

We first measured dwell times for β:T238A tubulin in GMPCPP (Fig. 4B). Like wild-type, β:T238A tubulin dwell times spanned five orders of magnitude, including dwells exceeding 100 s. A tri-exponential fit was required to fit the dwell time distribution, yielding characteristic dwell times of 137 ms (fast, 73%), 2.6 s (intermediate, 22%), and 67 s (slow, 5%). The characteristic dwell times were 2-5-fold longer for β:T238A tubulin than wild-type, indicating that the mutation modestly increases the tubulin:microtubule affinity in GMPCPP.

The β:T238A mutation primarily affects the microtubule shrinking rate (Geyer et al., 2015; Ye et al., 2020), which is a property of the GDP-lattice. Accordingly, we measured β:T238A dwell times on microtubule ends in GDP and found much more substantial differences to wild type than in GMPCPP. Most strikingly, at both ends of the microtubule the distribution of β:T238A dwell times in GDP showed the three-phase binding that had been lost for wild-type in GDP (see above) (Fig. S4). The characteristic plus-end dwell times for β:T238A in GDP were 67 ms (fast, 76%), 556 ms (intermediate, 16%), and 14 s (slow, 8%). The fast and intermediate events had similar duration as wild-type in GMPCPP and were about two-fold longer than wild-type in GDP, indicating that β:T238A tubulin modestly stabilizes longitudinal interactions in GDP. By contrast, the β:T238A mutation had an outsized effect on the corner interactions: whereas the corresponding long dwells were reduced at least 100-fold between GMPCPP and GDP for wild-type, the reduction was only ∼8-fold for β:T238A. Furthermore, the slow β:T238A dwell time in GDP (14 s) is within 3-fold of the slow wild-type dwell time in GMPCPP (31 s). Thus, the buried β:T238A mutation selectively and substantially attenuates the sensitivity of corner interactions to nucleotide state.

## DISCUSSION

The experiments described here took advantage of an improved single-molecule assay that provides more faithful estimates of biochemical off-rates than its predecessor (Mickolajczyk et al., 2019b). The resulting data revealed a new state/transition on the microtubule end, clarified how nucleotide state influences microtubule stability, and demonstrated that the allosteric response to GDP can be tuned by amino acid substitutions in tubulin (Fig. 5).

**Fig. 5.**
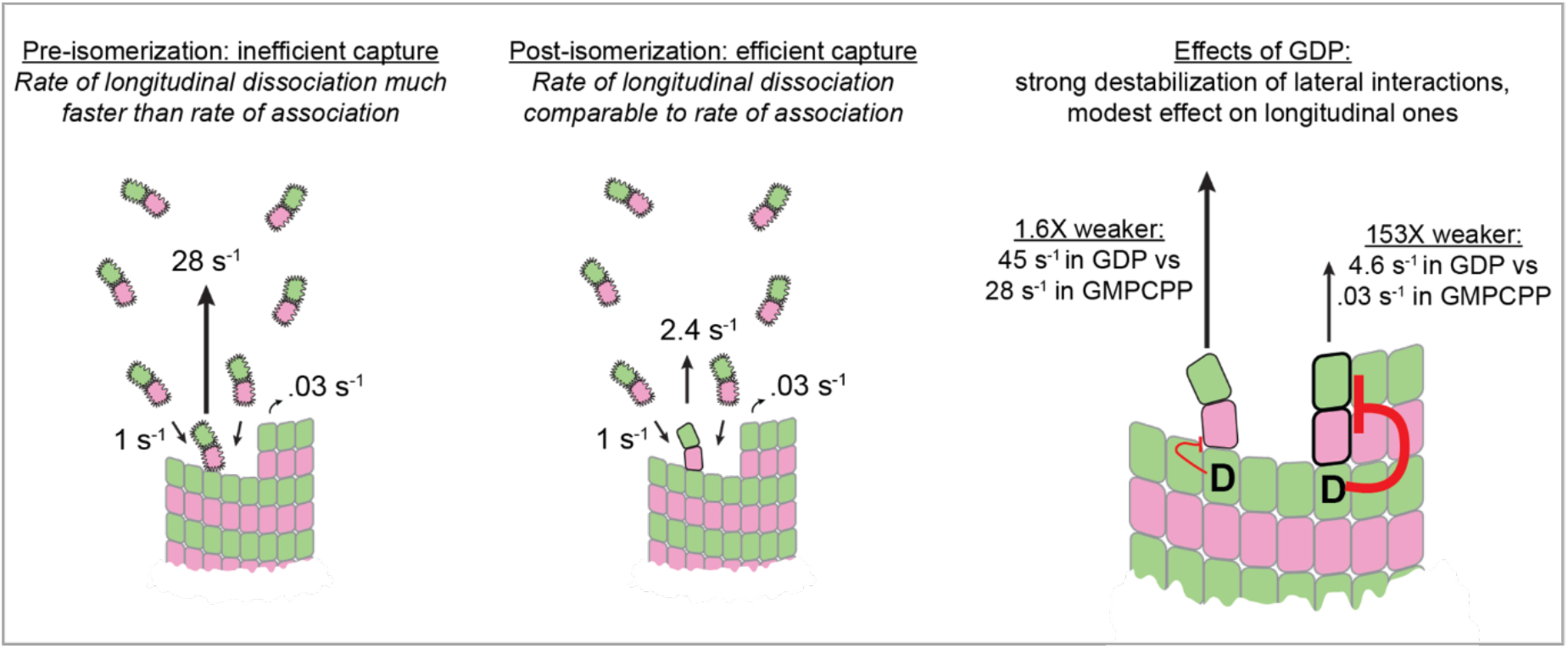
Summary of results and insight into microtubule growth dynamics. (Left) Tubulin dimers initially make weak longitudinal interactions that dissociate faster than they can be stabilized by association of a neighboring tubulin. (Middle) Isomerization strengthens the longitudinal contact such that dissociation and association rates become comparable, which makes stabilization by association of a neighboring tubulin much more likely. (Right) GDP at the tubulin:tubulin interface acts allosterically to weaken lateral interactions by >100-fold, with only a ∼2-fold effect on longitudinal interactions.

### Microtubules promote their own growth through a previously undetected ‘maturation’ process that provides stronger longitudinal contacts with the microtubule end

Our single-molecule measurements of tubulin dwell times on the microtubule end unexpectedly revealed three classes of binding site that we ascribed to two forms of longitudinal and a corner interaction. This previously undetected state provides evidence for a kind of positive feedback that allows microtubules to privilege their own elongation by promoting a longer-lived form of longitudinal interaction that is more likely to be stably incorporated into the microtubule (Fig. 5, middle).

What kind of change underlies the isomerization to the stronger form of longitudinal interaction? Based on recent evidence that curved protofilament extensions are present at the end of growing microtubules (Atherton et al., 2017; McIntosh et al., 2018; van den Berg et al., 2023), and for reasons explained in the next section, the isomerization seems unlikely to require substantial ‘straightening’ of the newly-bound tubulin. Instead, we speculate that the isomerization reflects a more local ‘settling in’ conformational change like loop or other movements that reflect an adaptation at the tubulin:microtubule interface. This view is reminiscent of a conformational adaptation process that has been proposed to occur during the polymerization of FtsZ, a distantly related bacterial ortholog of tubulin that forms single-stranded filaments stabilized solely by longitudinal interactions (Erickson, 2019; Miraldi et al., 2008).

The idea that microtubules reinforce their own elongation by promoting a stronger form of longitudinal tubulin:tubulin interaction has potential implications for spontaneous and templated microtubule nucleation (Kollman et al., 2011; Roostalu & Surrey, 2017). For spontaneous nucleation, if longitudinal interactions between unpolymerized tubulins are less able to promote the ‘matured’ state, then the formation of small tubulin oligomers would be selectively disadvantaged, representing an additional barrier to spontaneous nucleation. Templated nucleation by γ-Tubulin Ring Complexes (γTuRCs) involves αβ:γ interactions (Kollman et al., 2011; Roostalu & Surrey, 2017), so the degree to which longitudinal interactions with γ-tubulin also promote the isomerization of αβ-tubulin to a more tightly bound state should strongly influence the earliest steps in templated nucleation.

### GDP selectively weakens lateral tubulin:tubulin interactions, with little effect on longitudinal ones

GTP hydrolysis destabilizes the microtubule lattice, but in the absence of biochemical data there have been conflicting ideas about how it does so. Our measurements indicate that corner interactions are much (>100-fold) weaker in GDP compared to GMPCPP even though longitudinal interactions are only modestly affected (2-fold weaker). Corner affinity reflects the combination of longitudinal and lateral interactions, so a strong effect on corner affinity combined with a weak effect on longitudinal affinity indicates that GDP selectively weakens lateral contacts (Fig. 5, right).

Lateral interactions between αβ-tubulins require at least partial ‘straightening’ (Brouhard & Rice, 2014, 2018; Ravelli et al., 2004; Wang & Nogales, 2005). Our finding that GDP acts at a distance to destabilize lateral/corner interactions is most simply explained if GDP less effectively promotes/stabilizes straighter conformations of αβ-tubulin on the microtubule end compared to GTP. Our data do not distinguish between two possible models: that GDP makes the process of straightening harder (akin to stiffening a spring) or that GDP makes lateral contacts weaker and therefore less able to maintain the straight conformation (akin to weakening a latch). Either way, because the nucleotide resides at the longitudinal tubulin:tubulin interface, ‘stiffening’ tubulin or weakening the lateral latch both imply that the straight conformation of αβ-tubulin is stabilized at the microtubule end by a long-range, allosteric response to GTP. We recently showed that GDP exposure on the microtubule end poisoned elongation (McCormick et al., 2024); our new results suggest that the predominant mechanism of GDP-induced poisoning is to selectively destabilize the high-affinity corner interactions that are required for microtubule growth.

### The strength of tubulin’s allosteric response to GDP can be tuned by mutations

Our measurements of microtubule end residence times for the microtubule-stabilizing β:T238A mutant provides further insight into the role of allostery in microtubule dynamics and how it can be tuned by amino acid substitutions. In GDP, the β:T238A mutation restored (to within 5-fold) the corner interactions that were lost in wild-type tubulin. Thus, the mutation attenuates the weakening of lateral interactions that normally accompanies GTP hydrolysis and that underlies depolymerization. In previous work the β:T238A mutation did not detectably affect conformational strain energy in depolymerizing yeast microtubules (GDP state) (Driver et al., 2017), which suggests that the mutation strengthens the lateral latch rather than softening the curved tubulin ‘spring’.

### Concluding remarks

This work provides multiple insights into the biochemical mechanisms underlying microtubule dynamics: it reveals a new transition through which microtubules promote their own elongation, it clarifies how GDP affects the strength of lattice contacts, and it demonstrates that the allosteric response to nucleotide state can be tuned by amino acid variation. These findings raise new questions about the microtubule cytoskeleton more broadly. Do any regulatory factors act on the isomerization process to promote or inhibit the transition to the more strongly bound longitudinal interaction? If a relatively conservative mutation like Threonine to Alanine can so dramatically affect the strength of tubulin: microtubule interactions, is the scope for species-or isotype-specific variation in microtubule stability and dynamics greater than revealed to date? We expect that answers to these and other questions will come from future applications of the improved single-molecule assay presented here.

## METHODS

### Recombinant human tubulin expression and purification

Plasmids for expressing recombinant human tubulin were designed in pFastBacDual (Thermo Fisher Scientific) with an internal His_6_-tag on α1b tubulin (TUBA1B) and a C-terminal Sortase A recognition sequence and Strep-tag II on β3 tubulin (TUBB3). The β:T238A mutant of TUBB3-tubulin was generated using QuikChange (Agilent) mutagenesis, using the wild-type TUBB3 expression plasmid as the template and primers designed according to the manufacturer’s instructions. Recombinant baculoviruses were produced using the Bac-to-Bac System (Thermo Fisher Scientific) and used to infect Tni insect cells (Expression Systems) at a density of 2 × 106 cells/ml. Wild-type and mutant human αIb/β3-tubulin were purified through sequential nickel and strep-tactin affinity chromatography.

Biotin labeling of the sortase epitope containing tubulin purified as described above was performed employing a two-step method. First, we prepared a biotinylated peptide by mixing equal volumes of a GGGC peptide (5 mM in 25 mM PIPES pH 7.0) (Genscript) with EZ-Link Maleimide-PEG2-Biotin (MPB; 100 mM in DMSO) (Thermo Fisher Scientific) and allowed them to react for two hours at room temperature before quenching with 100 mM DTT for 30 minutes at room temperature. Second, the biotinylated peptide was ligated to tubulin-LPETGG using sortase in a reaction containing 6 µM human tubulin, 120 µM GGGC-MPB, 30 mM DTT, 50 µM GTP, and 2.5 µM Sortase A in sortase buffer (20 mM Tris pH 7.5, 125 mM NaCl, 10 mM CaCl_2_, and 100 µM TCEP). This procedure also removes the Strep-tag II. To separate tubulin from unreacted peptide and sortase, the reaction products were further purified over a Source Q column and exchanged into BRB80 (80 mM K-PIPES pH 6.9, 1 mM MgCl_2_, 1 mM EGTA, 50 μM GTP) using 7K MWCO Zeba Spin Desalting Columns (Thermo Fisher Scientific), flash-frozen, and stored at −80 °C. The extent of biotin labeling and removal of source components was assessed by whole protein mass spectrometry at the UT Southwestern Proteomics Core facility.

### Nano-gold-labeling

On the day of the experiment, biotinylated wild-type or β:T238A tubulin was thawed, diluted, and centrifuged in a Beckman Airfuge at 199,000 x g (30 psi) for 10 minutes to remove aggregates. The tubulin concentration (∼ 1 μM) was determined by absorbance at 280 nm using ε_tubulin_ of 115,000 M^−1^ cm^−1^. The stock concentration of 20 nm gold nanoparticles conjugated to streptavidin (BBI Solutions) was determined using iSCAT microscopy by counting individual particles bound to a surface uniformly coated with 1 mg/mL biotinylated tubulin. Based on the analysis, the concentration of the stock solution was estimated to be 2.8 nM, which is slightly lower than the 4 nM value specified by the manufacturer. Next, 2.5 nM gold-streptavidin conjugates were mixed with the 80-100 nM biotinylated tubulin (wild-type 27 % and T238A mutant 38% biotinylated) at a gold:tubulin ratio of 1:10 (considering the extent of biotin labeling) and were incubated for 20 minutes. The 1:10 gold-to-tubulin ratio was used to maximize the number of interactions of gold-tubulin with microtubule ends, and a 1:1 ratio showed similar landing behavior with far fewer counts.

### Purification and DNP labeling of bovine tubulin

PC-grade bovine brain tubulin was purified as previously described (Cleary et al., 2022; McCormick et al., 2024), cycled twice through polymerization/depolymerization, quantified by absorbance at 280 nm using ε_tubulin_ of 115,000 M^−1^ cm^−1^, and diluted to 80 µM in BRB80. Samples were aliquoted, flash frozen in liquid nitrogen, and stored at –80 °C. Before experiments, tubulin aliquots were thawed on ice, diluted to 20 µM in BRB80, and concentrations reconfirmed by A280.

For labeling, bovine tubulin was conjugated with EZ-Link 2,4-dinitrophenyl NHS-ester (DNP-NHS) following the protocol provided by the manufacturer (Avantor). Briefly, microtubules were polymerized in BRB80 from a solution of 40 μM tubulin, 1 mM GTP, 4 mM MgCl_2_, and 5% DMSO for 45 min at 37 °C. Next, an equimolar amount of DNP-NHS dissolved in DMSO (ThermoFisher 20217) was added and reacted for 35 min at 37 °C. The labeled microtubules were pelleted by airfuge at 30 psi for 10 minutes and the pellet was resuspended in cold BRB80 and depolymerized by incubating on ice for 45 min. The tubulin solution was airfuged again for 10 min at 30 psi to remove any remaining polymerized tubulin, and the supernatant containing DNP-labeled tubulin was collected. The concentration of labeled tubulin was quantified by A280 nm using ε_tubulin_ of 115,000 M^−1^ cm^−1^, and aliquoted samples frozen on liquid nitrogen for later use.

### DNP-labeled GMPCPP-microtubule seeds

To generate DNP-labeled microtubule seeds for gold-tubulin binding experiments, 20 μM DNP-labeled tubulin was mixed with 1 mM GMPCPP and 1 mM MgCl2, then incubated at 37 °C for 60 minutes to nucleate microtubule seeds. The solution was then diluted 1:2 with BRB80 containing 1 mM GMPCPP and 1 mM MgCl2 and incubated at 37 °C for an additional 2 hours to extend the seeds. The microtubules were pelleted by airfuging at 30 psi for 10 minutes, resuspended in BRB80 with 20% glycerol, aliquoted into smaller fractions, flash-frozen in liquid nitrogen, and stored at –80 °C. Before experiments, an aliquot was thawed at 37 °C, diluted and the microtubules pelleted to remove glycerol. The pellet was resuspended in BRB80 supplemented with 1 mM GMPCPP and 1 mM MgCl2, then diluted as needed for assays.

### iSCAT microscopy

Imaging was carried out on a custom-built iSCAT microscope, constructed as previously described (Mickolajczyk et al., 2019) with minor modifications. The instrument was built around a Mad City Labs RM20 stage and a 60x, 1.49 numerical aperture objective (Nikon) equipped with an objective heater (HT10K, Thorlabs). A 520 nm laser (OFL420-1000-TTL; OdicForce) was scanned in two dimensions across the sample using an acousto-optic deflector (DRFA10Yxx; AA Opto-Electronic). The 2D laser scanning pattern was controlled using two function generators producing sawtooth waveforms that were synchronized using a trigger signal from the camera. Images were taken using a Photron Fastcam Nova S16 (Photron) controlled by custom-written LabVIEW software. The camera exposure time was 1 ms, and the laser was scanned over the sample 200 times per exposure to achieve uniform illumination. Movies were captured at 125 frames/s and initially saved to a high-speed drive using FASTDock; they were subsequently transferred to a computer for further analysis using ImageJ. The total microscope magnification was 420×, resulting in an image calibration of 46.7 nm per pixel.

### Reversible tubulin binding assay and polarity marking

Coverslips (1½, 24 × 30 mm^2^; Corning) were cleaned by gentle heating for 2-3 hours in detergent solution (ICN Biomedicals, CAT No: 76-670941), thoroughly rinsed with ddH_2_O, and cleaned for 12 minutes in a plasma cleaner (Harrick Plasma). Silanization was performed using 1H,1H,2H,2H-perfluorodecyltrichlorosilane (Alfa Aesar L165804-03), and the hydrophobicity was confirmed with a droplet test. The silanized coverslips were fixed onto microscope slides using double-sided tape to construct 10 μL flow cells. A 4 mg/ml solution of DNP-labeled antibody (Sigma-Aldrich, D9656) was flowed into the chamber, followed by a blocking step with 5% Pluronic F127 and 2 mg/mL casein to prevent non-specific gold nanoparticle binding to the glass surface. DNP-labeled microtubule seeds were introduced into the flow cell, incubated for 5 minutes to allow for surface attachment through the antibody, and unbound MT seeds were washed out. Finally, a solution containing 1 nM gold-tubulin, 1 mM GMPCPP, 1 mM MgCl_2_, 0.08 % methylcellulose in BRB80 was introduced to the flow cell. In some experiments unlabeled tubulin (0.25 or 1.0 μM final concentration) was added to enable microtubule growth. In all experiments, temperature was kept at 30 °C using an objective heater. Movies were recorded at 125 frames per second over 3-4 minutes (25,000-30,000 frames). Multiple movies were recorded consecutively from the same flow cell and no flow cell was imaged for longer than 25 minutes. Experiments were repeated using different flow cells, tubulin purified from different batches, and different stocks of gold-streptavidin conjugate. For the tubulin binding assay in 1 mM GDP, the gold-tubulin and the immobilized microtubule seeds were pre-incubated in 1 mM GDP for 10 minutes to allow nucleotide to exchange into the free tubulin and the exchangeable site at the terminal tubulin on the plus-end of the seed. Mutant tubulin experiments were carried out in similar conditions to the wild type.

After acquiring binding events, microtubule polarity was determined as follows. After washing out the gold-tubulin solution, a polymerization mix was added that contained 20 μM unlabeled tubulin and 10 μM NEM-labeled tubulin to suppress minus end growth, 1 mM GTP, 1 mM MgCl_2_, and 0.08% methylcellulose. Movies of growing microtubules were recorded for 5–7 minutes at 2 frames per second and the faster growing end was assigned as the plus-end. Notably, with this NEM–tubulin procedure, over 90% of surface-attached microtubules showed no detectable minus-end growth.

### Image processing and data analysis

Each movie typically contained 40–70 stabilized microtubules and typically yielded 5–9 binding events. Because the camera recorded at 125 frames per second, durations were discretized in 8 ms intervals, and the shortest measurable binding duration was 16 ms. An event was classified as irreversible if gold-tubulin landed on a microtubule tip and remained bound for the flow cell’s lifetime (longer than 200 seconds). For each condition, dwell times of reversible events were plotted as a cumulative distribution function (CDF) using one point per discretized frame time. To account for missed events, the data were fit using a lower cutoff (see equations S1 and S2). Due to discretization of time, there was no simple way to objectively set the minimum cutoff duration. We fit multiple data sets either using fixed values or setting the lower cutoff as a free parameter and found that the resulting fit parameters were highly sensitive to the cutoff choice. Thus, we chose the lower cutoff value empirically by fitting multiple data sets while setting the lower cutoff to a free parameter, taking the average cutoff value from these fits, and adopting a consensus lower cutoff of 19 ms for all subsequent fits.

We fit the cumulative distribution functions (CDFs) of the measured dwell times using MATLAB, with either a bi-exponential or tri-exponential function, as follows:

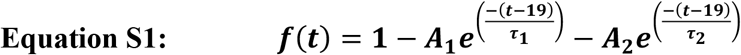

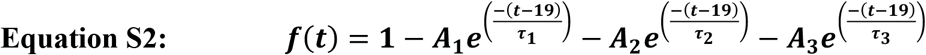

In the fits, τ_1_, τ_2_ and τ_3_ represent the characteristic dwell times and A_1_, A_2_ and A_3_ represent relative amplitudes. The amplitudes were corrected based on the lower cutoff, t_0_ = 19 ms by the following equation:

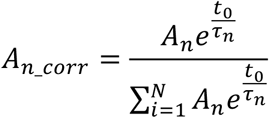

As shown in Fig. 1D (inset), the bi-exponential function fails to adequately capture the long dwells in GMPCPP. The inability of a bi-exponential function to fit the observed dwell times in GMPCPP is a robust result: it persists over a range of different starting values for the fitted parameters. Moreover, when we manually force the bi-exponential fit to better match the long dwell times, the fit to the short dwell times became substantially worse, further supporting the inadequacy of the bi-exponential model and the need for a tri-exponential description. The Best-fit parameters determined from the bi-or tri-exponential function are reported as the fit value ± 95 % confidence interval (CI). Since dwell times or event fractions cannot be negative, any negative lower bounds were adjusted to 0.00.

Since all minus ends were not unambiguously growing at the free tubulin concentrations used, we did not analyze the minus end tubulin-binding durations in the presence of 0.25 and 1 uM unlabeled tubulin.

### Computational modeling

We performed kinetic Monte Carlo simulations of growing MT plus-ends using the simulation code and analysis algorithms described previously (Kim & Rice, 2019; Mickolajczyk et al., 2019; Piedra et al., 2016) with minor modifications. In this method, the MT lattice is represented as a 2D array with a periodic boundary condition to mimic the cylindrical wall of the MT, and microtubule dynamics are simulated one biochemical reaction at a time. Biochemical reactions are chosen at random in accordance with their relative probabilities, which is determined by the associated rate constants. For the simulations shown in this work, the possible reactions are: (i) αβ-tubulin association at the end of a protofilament (rate: k_on_[αβ-tubulin], where k_on_ was determined as described in Fig. 2), (ii) αβ-tubulin dissociation from the end of a protofilament (three possible rates depending on the nature of the site: weak longitudinal, ‘matured’ stronger longitudinal, and corner; rates were either taken as the inverse of the characteristic dwell times or calculated as described below), and (iii) ‘isomerization’ from weak to more strongly-bound longitudinal site. We did not include GTPase because all experiments used GMPCPP, a hydrolysis-resistant GTP analog. The isomerization reaction was modeled as an irreversible first-order process. Simulations result in a biochemical ‘movie’ of microtubule growing or shrinking, the rates of which can be calculated from simulated microtubule length vs time.

The presence of an isomerization reaction means that there are two different ways for a subunit to ‘leave’ the weakly bound longitudinal state: dissociation or isomerization to the matured state. Consequently, the short characteristic dwell time we measured reflects contributions from both dissociation and isomerization: τ_short_ = 1/(k_offWeak_+k_iso_). Because we measured τ_short_=0.0346 s, we set k_offWeak_+k_iso_ = 28.8 s^-1^. How to determine the values for k_offWeak_ and k_iso_? Considering only fast and intermediate events as the possible events, then the proportion of fast events provides a second constraint. The proportion of fast events is given by k_off_^Weak^/ (k_off_^Weak^ + k_iso_) and can be calculated from the measured amplitudes of each kind of event (Fig. 1D): proportion of fast events = (51% fast) / (51% fast+43% intermediate) = 0.543 = 1/1.84. Using the two relationships k_off_^Weak^+k_iso_ = 28.8 s^-1^ and k_off_^Weak^/(k_off_^Weak^+ k_iso_)=1/1.84 yields k_off_^Weak^=15.65 s^-1^ and k_iso_=13.15 s^-1^. Likewise, the off-rate for the post-isomerized state should reflect the time spent in the pre-isomerized state plus the lifetime of isomerized state, in other words the measured characteristic time of 0.411 s corresponds to 0.034 s + 1/k_off_ ^intermediate^, which yields k_off_ ^intermediate^= 2.65 s^-1^. Finally, the characteristic dwell time for corner interactions, 30.6 s, is so much longer than the short or intermediate dwells that their contribution can be neglected, so k_off_ ^Corner^ = 1/30.6 s^-1^ = 0.033 s^-1^.

## ACKNOWLEDGMENTS

This study was supported by National Institute of Health Grants NIH R35 GM139568 to W.O.H, and NIH R01 GM135565 and R35 GM156385 to L.M.R. We thank Daguan Nong for iSCAT microscopy reconstruction and Joseph Cleary for early experimental efforts. We also thank all members of the W.O.H. and L.M.R. laboratories for helpful discussions.

## Author contributions

E.B. purified recombinant proteins and performed site specific biotinylation, S.M. purified bovine tubulin, performed all single molecule assays, and analyzied data, L.M.R performed the computational simulations. W.O.H. and L.M.R conceived ideas, designed research, discussed the concept throughout. S.M, W.O.H and L.M.R wrote and all authors edited the manuscript.

## Competing interests

The authors declare no competing interests.

Correspondence and requests for materials should be addressed to Will Hancock or Luke Rice

## Data availability

Source data will be provided with the paper.

## Code availability

Computational simulations of microtubule dynamics relied on lightly modified versions of previously described codes (Cleary et al 2022 and McCormick et al 2024) that are publicly available (https://git.biohpc.swmed.edu/s422146/simulate-mt-44 and https://git.biohpc.swmed.edu/ricelab/simulate-mt-52)

