## Supplemental Material for "Mechanisms of microtubule dynamics from single-molecule measurements"

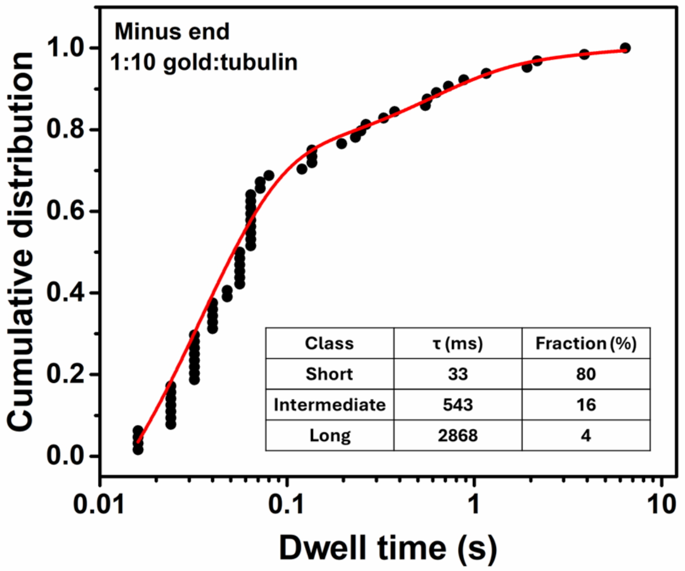


**Fig. S1. Distribution of minus-end gold-tubulin dwell times in GMPCPP reveals three classes of binding.** Cumulative distribution function (CDF) for the measured dwell times (circles); n = 64 dwell events. CDF was fit by a tri-exponential function (red line) with characteristic dwell times 33 [26-41] ms for 80 % of dwells, 543 [0-150] ms for 16 % of dwells, and 2.9 [0-19] s for 4 % of dwells. Compared to plus-ends (Fig. 1D), minus ends exhibit a higher proportion of short-lived binding events and a reduced duration of the long-lived binding events. Both of these differences in principle could contribute to the slower minus-end growth rate.

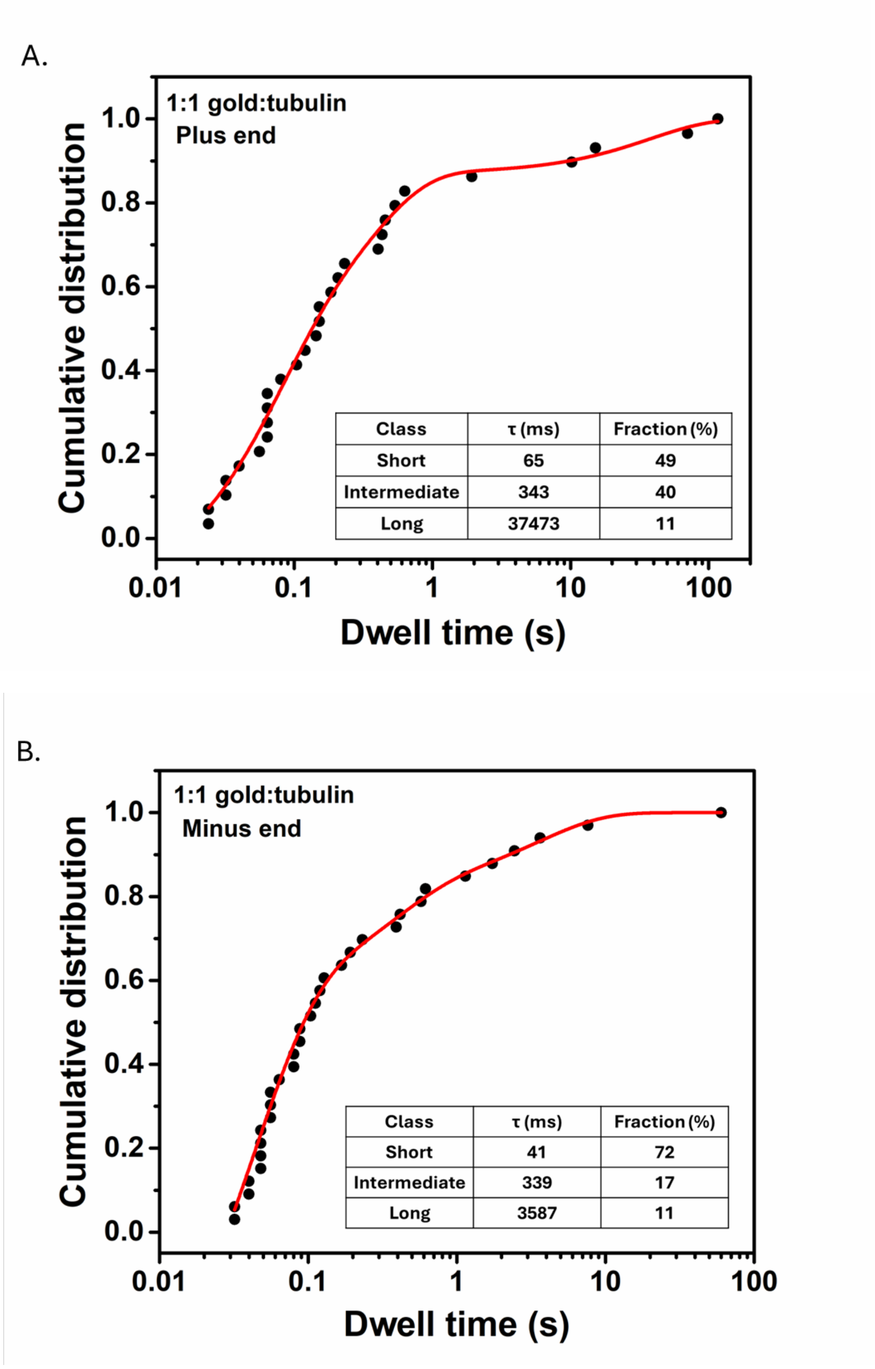


**Fig. S2. Three characteristic binding durations were also observed at both ends using a gold:tubulin ratio of 1:1.** A. Cumulative distribution function (CDF) for the measured plus-end dwell times (circles, n = 40) with the tri-exponential fit (red line). The best fit characteristic times at the plus end were 65 [14—120] ms for 49 % of all dwells, 343 [91—600] ms for 40 % of all dwells, and 37 [0.00—88] s for 11 % of all dwells. **B.** Minus end dwell times with best fit characteristic times 41 [31—51] ms for 72 % of all dwells, 339 [69—610] ms for 17 % of all dwells, and 3.6 [1.0—6.7] s for 11% of all dwells (n = 33). This control experiment argues against the possibility that intermediate binding durations result from multivalent tubulin-microtubule interactions.

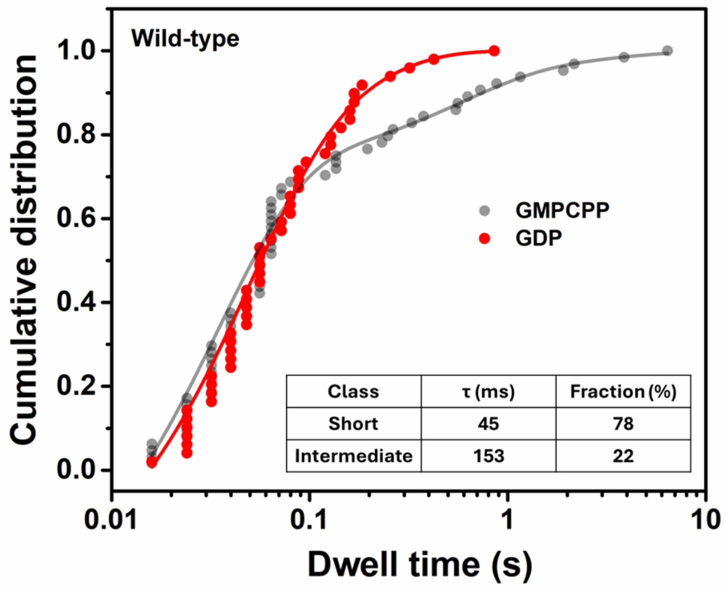


**Fig. S3. Minus end dwell time distributions for wild type tubulin in GDP.** CDF of dwell times (red circles; n = 56 events) and bi-exponential fit (red line) with characteristic times 45 [27—64] ms for 78 % of all dwells and 153 [0.00—311] ms for 22 % of all dwells. In GDP two binding phases were observed at the minus end with no long dwells detected (inset), consistent with the plus-end results. Minus end dwell time distributions in GMPCPP (in gray, reproduced from Fig. S1) are shown for comparison. The absence of long dwells in GDP at both ends reinforces the idea that GDP-bound tubulin is unable to form lateral interactions.

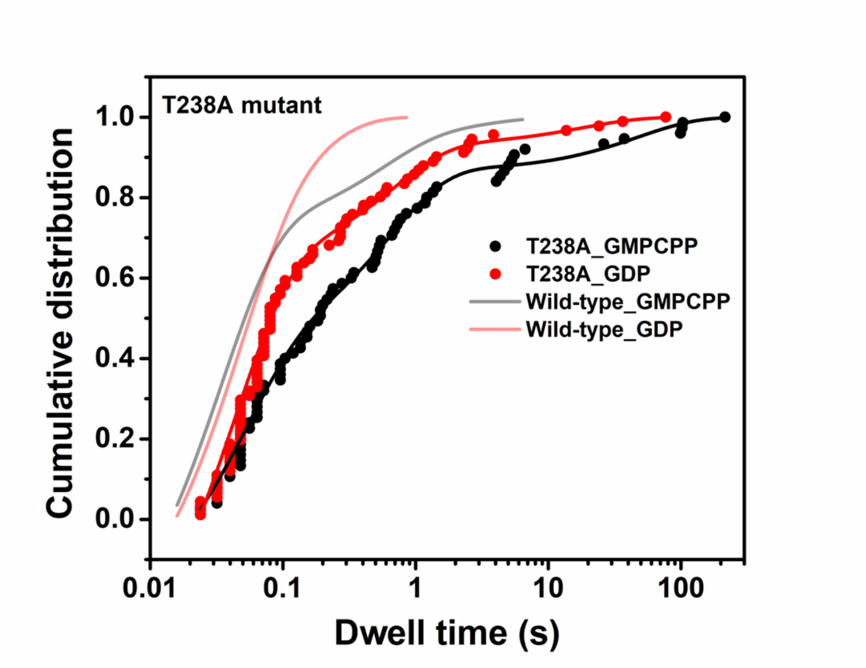


| | Fast | Intermediate | Slow |
|---|---|---|---|
| WT_GMPCPP | 33 ms (0.8) | 541 ms (0.16) | 2.9 s (0.04) |
| WT_GDP | 45 ms (0.82) | 185 ms (0.18) | **** |
| T238A_GMPCPP | 48 ms (0.53) | 634 ms (0.36) | 48 s (0.1) |
| T238A_GDP | 42 ms (0.74) | 659 ms (0.21) | 18 s (0.05) |

**Fig. S4. Minus end dwell time distributions for β:T238A mutant tubulin in GMPCPP and GDP.** CDF for T238A-tubulin in GMPCPP (black) and GDP (red) fit with tri-exponential functions. Light red and black curves are fits to for wild type minus end durations in GDP and GMPCPP (reproduced from Fig. S3). The characteristic times for the mutant in GMPCPP were 48 [40-57] ms for 53 % of all dwells, 634 [550-720] ms for 36 % of all dwells, and 48 [27-69] s for 10% of all dwells. The characteristic times for the mutant in GDP were 42 [37-47] ms for 74 % of all dwells, 659 [480-840] ms for 21 % of all dwells, and 18 [0-37] s for 5% of all dwells. n =75 and 91 dwell events, respectively. The table summarizes the binding events, categorized as fast, intermediate, and slow phases, with corresponding dwell time and fraction of contributions (in parentheses). Notably, compared to wild-type tubulin in GMPCPP, the mutant showed a higher proportion of intermediate events and a 15-fold stabilization of long dwells, suggesting that the mutation strengthens both intermediate and long binding interactions. In GDP, the mutant retained three binding phases, like the plus end (Fig. 4), and displayed a strengthened intermediate phase, further supporting the idea that the T238A mutation allosterically stabilizes tubulin-tubulin interactions independent of the nucleotide state.

**Description of Additional Supplementary Files**

**Movie S1.**
Example movie of a fast (40 ms) reversible gold-tubulin binding event at the microtubule end (125 frames per second).
Movie speed 4 frames per second.
Scale bar 1 μm.

**Movie S2.**
Example movie of an intermediate (440 ms) reversible gold-tubulin binding event at the microtubule end (125 frames per second).
Movie speed 15 frames per second).
Scale bar 1 μm.

**Movie S3**
Example movie of a slow (25 s) reversible gold-tubulin binding event at the microtubule end (125 frames per second).
Movie speed 500 frames per second.
Scale bar 1 μm.

**Movie S4**
Example movie of an irreversible gold-Tubulin binding event (125 frames per second). The microtubule shows thermal fluctuations, and the gold-tubulin fluctuates with it.
Movie speed 1000 frames per second).
Scale bar 1 μm.

**Movie S5**
Example movie of a reversible gold-Tubulin binding event (125 frames per second) at both ends of a microtubule.
Movie speed 7 frames per second).
Scale bar 1 μm.
